# Invasion genetics of the longhorn crazy ant: the global expansion of a double-clonal reproduction system

**DOI:** 10.1101/2022.03.17.484689

**Authors:** Shu-Ping Tseng, Hugo Darras, Po-Wei Hsu, Tsuyoshi Yoshimura, Chow-Yang Lee, James K. Wetterer, Laurent Keller, Chin-Cheng Scotty Yang

## Abstract

Reproduction mode represents a key determinant for success of biological invasion as it influences the genetic variation and evolutionary potential of introduced populations. The world’s most widespread invasive ant, *Paratrechina longicornis*, was found to display an unusual double-clonal reproduction system, whereby both males and queens were produced clonally, while workers are produced sexually. Despite its worldwide distribution, the origin of this ant species and the prevalence of the double-clonal reproductive system across the ant’s geographic range remain unknown. To retrace the evolutionary history of this global invasive species and its reproductive system, we examined genetic variation and characterized the mode of reproduction of *P. longicornis* sampled worldwide using both microsatellite genotyping and mitochondrial DNA sequencing approaches. Analyses of global genetic variations indicate that the Indian subcontinent is a genetic diversity hotspot of this species, suggesting that this geographic area is at least part of its native range. Our analyses revealed that inferred native and introduced populations both exhibit double-clonal reproduction. Remarkably, queens and males worldwide belong to two separate, non-recombining clonal lineages. Workers are highly heterozygous and first-generation inter-lineage hybrids, a pattern strongly supportive of a strict worldwide prevalence of double clonality. By maintaining heterozygosity in the worker force, this unusual genetic system allows *P. longicornis* to avoid inbreeding during colonization bottlenecks and may have acted as an adaptive trait linked to the species’ invasion success.

## INTRODUCTION

Globalization and commerce have significantly facilitated the spread of exotic species and increased the threat of new biological invasions (Bertelsmeier et al., 2017; Early et al., 2016; Meyerson and Mooney, 2007; Occhipinti-Ambrogi and Savini, 2003). Social insects are among the most successful invaders and account for half of the worst invasive insect species listed by the IUCN’s Global Invasive Species Database (IUCN: SSC Invasive Species Specialist Group, n.d.). Despite their undoubted success as invasive species, most social Hymenopterans have a sex-determination system that would seem to run counter to their capacities for invasiveness, namely single-locus complementary sex determination (sl-CSD) (Schmieder et al., 2012). Under sl-CSD, a queen mated with a male sharing one of her alleles at the sex determination locus produces diploid progeny of which half are males (Cook and Crozier, 1995; Crozier, 1977, 1971; Ross and Fletcher, 1985; Stouthamer et al., 1992). Production of diploid males represents a cost for colonies because they are typically inviable or infertile (Cowan and Stahlhut, 2004; Heimpel and de Boer, 2008; Naito and Suzuki, 1991; Wilgenburg et al., 2006; Zayed and Packer, 2005). In small founding populations with low genetic diversity, diploid male production can be costly and potentially result in population extinction (Hedrick et al., 2006; Ross and Fletcher, 1986; Whitehorn et al., 2009; Wilgenburg et al., 2006; Zayed and Packer, 2005). Some forms of asexual reproduction may also help circumvent the disadvantages associated with inbreeding (Hagan and Gloag, 2021; Rabeling and Kronauer, 2013).

Reconstructing species invasion histories is crucial, especially since such information can benefit the implementation of control measures against these invaders, uncover natural biocontrol agents, and identify characteristics that predispose invasive species to thrive in new environments (Colautti et al., 2004; Jeschke, 2014; Sakai et al., 2001; Wilson et al., 2009). For example, factors contributing to the invasion success could be deduced from comparisons of traits such as reproductive strategies and colony structures between invasive and native populations. Since (Lee and Yang, 2022; Wetterer, 2010, 2009, 2005).

The longhorn crazy ant, *Paratrechina longicornis* (Latreille, 1802), is a common household, garden, and agricultural pest (Wetterer, 2008). This species disrupts ecosystems by enhancing populations of phloem-feeding Hemipterans and displacing other invertebrates (Koch et al., 2011; Wetterer, 2008; Wetterer et al., 1999). It thrives in anthropogenically modified environments (LaPolla et al., 2013) and is frequently transported by cargo and other human-associated articles (Bertelsmeier et al., 2018; Lester, 2005; Weber, 1939). Consequently, *P. longicornis* has spread across most subtropical and tropical areas of the world for over a century and became arguably the most widely-distributed ant on Earth today (Fig. 1)(Wetterer, 2008). Despite the long-establishment history of *P. longicornis*, research on its invasion history has lagged behind other invasive ants, partly due to controversies over its native range. Attempts to deduce the geographic origin of *P. longicornis* from historical records, phylogenetic position, and habitat preferences failed to reach consensus (LaPolla et al., 2010; LaPolla and Fisher, 2014; Wetterer, 2008). *Paratrechina longicornis* has been recorded in undisturbed habitats in Southeast Asia (Wetterer, 2008) and is related to the genera *Pseudolasius* and *Euprenolepis*, which are both restricted to southeastern Asia (LaPolla et al., 2010), indicating that this ant may have a Southeast Asian origin. Yet, this Asian origin has been recently challenged by the finding of several congeneric species in Africa, suggesting that *P. longicornis* may be native to the African continent (LaPolla and Fisher, 2014). To date, no genetic data are available to untangle these hypotheses (Cristescu, 2015; Kirk et al., 2013; Lawson Handley et al., 2011). The reproductive biology of this species also remains puzzling. An unusual double-clonal reproductive system, whereby queens are clones of their mother, males are clones of their father, and sterile workers arise sexually has been previously reported in one population in Thailand (Pearcy et al., 2011). Whether this peculiar genetic system is widespread across the species range and could be related to the ant’s ecological success is unclear. Here, we investigate the population genetics, phylogeography, and breeding system of *P. longicornis* worldwide using both microsatellite and mitochondrial DNA (mtDNA) data for 256 localities sampled across most of the species’ range. These data allowed us to characterize the prevalence and distribution of double-clonality and resolve the question of the species’ native range, facilitating the reconstruction of the evolution of this global invasive ant species and its unique reproduction mode.

**Fig. 1.**
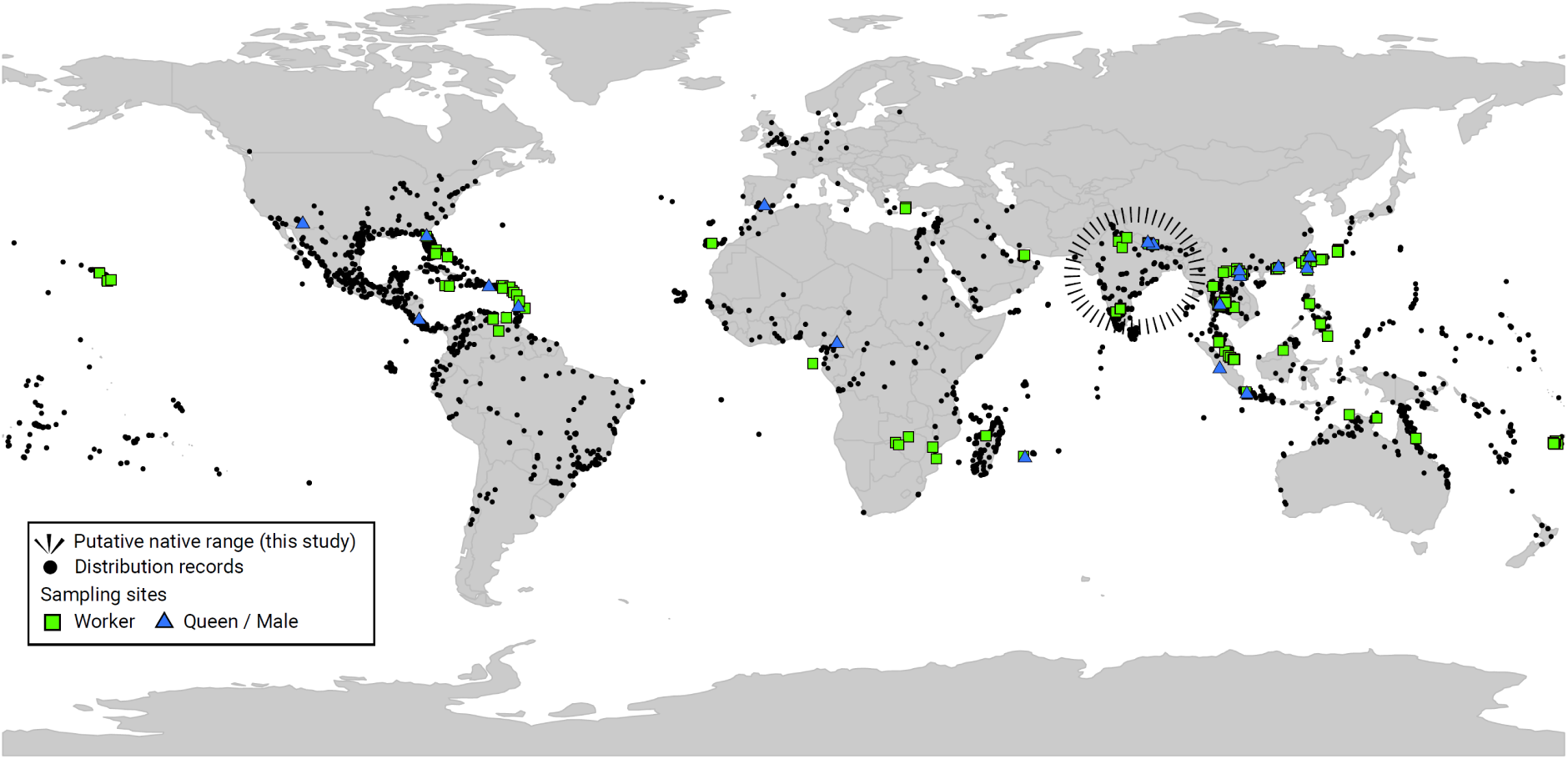
Distribution and sampling sites of *Paratrechina longicornis*. The 256 collection sites studied are indicated with squares and sites in which males or queens were collected are indicated by triangles (N = 22 localities). The global distribution of the species was obtained from Wetterer (2008), Antmaps.org, and iNaturalist.

## RESULTS

### Distribution of double-clonality

To investigate the distribution of double-clonality in *P. longicornis,* we analyzed the genotypes of workers, queens and males from 22 localities across Africa, America, Asia, and Europe (Table 1). Five males, which will be discussed later, had more than one allele at one or multiple markers and were excluded for the main comparative analyses. The genotypes of reproductive individuals from multiple localities clustered into two main groups (Fig. 2a). The first group contained all queen genotypes including the queen clone originally described in Thailand (Pearcy et al., 2011). The second group consisted of males only and included all the male genotypes inferred from queen- worker comparisons and the male clone identified by Pearcy et al. (2011) in Thailand. The two groups were distinguishable by their first PCoA coordinate and had diagnostic, non-overlapping allele size ranges at 19 of the 40 loci (Fig. 2b, S1, and S2). The observed high divergence between the two groups suggests that queens and males belong to two distinct lineages corresponding to the queen and male clonal genetic groups originally described in Thailand. Clustering analyses with STRUCTURE (Pritchard et al., 2000) confirmed these results. The second-order rate of change of the likelihood (Δ-K) was the highest for a predefined number of groups set to 2, and all runs performed with this number showed identical results with all individuals assigned to the same two respective lineages mentioned above with virtually no admixture (< 0.2% per individual; Fig. S3). The two lineages are hereafter referred to as Q and M lineages. All workers from the 22 localities were nearly completely heterozygous (mean individual heterozygosity ± SD = 91 ± 3%, range: 83-97%, N = 91 individuals) and carried one allele from each lineage at each of the 19 diagnostic loci. This suggests that workers were all Q/M hybrids produced from inter-lineage mating. To determine whether Q and M lineages identified in these 22 localities coexisted in all other populations, we investigated the allelic patterns of workers in the 234 localities where no reproductive individual was available. Workers were always extremely heterozygous (mean individual heterozygosity ± SD = 89 ± 4%, range: 75-98%; N= 250 workers; Fig. S4) with one allele typically corresponding to the size range of Q lineage and the other to the size range of M lineage at each of the 19 diagnostic loci (Fig. 2b and Fig. S1). Accordingly, in all but one locality (NP01) in Indian subcontinent, individual inbreeding coefficient across the 19 loci were lower than expected under random mating (mean Fij = -0,378; max = -0,073; N=251 workers, 95% confidence interval based on 1000 permutations of gene copies across individuals: [-0,016 to 0,014]). In line with these results, the PCoA coordinates of workers were all intermediate between those of the Q and M lineage groups (Fig. 2a). These results indicated that workers of *P. longicornis* are always first-generation hybrids of Q and M lineages.

**Fig. 2.**
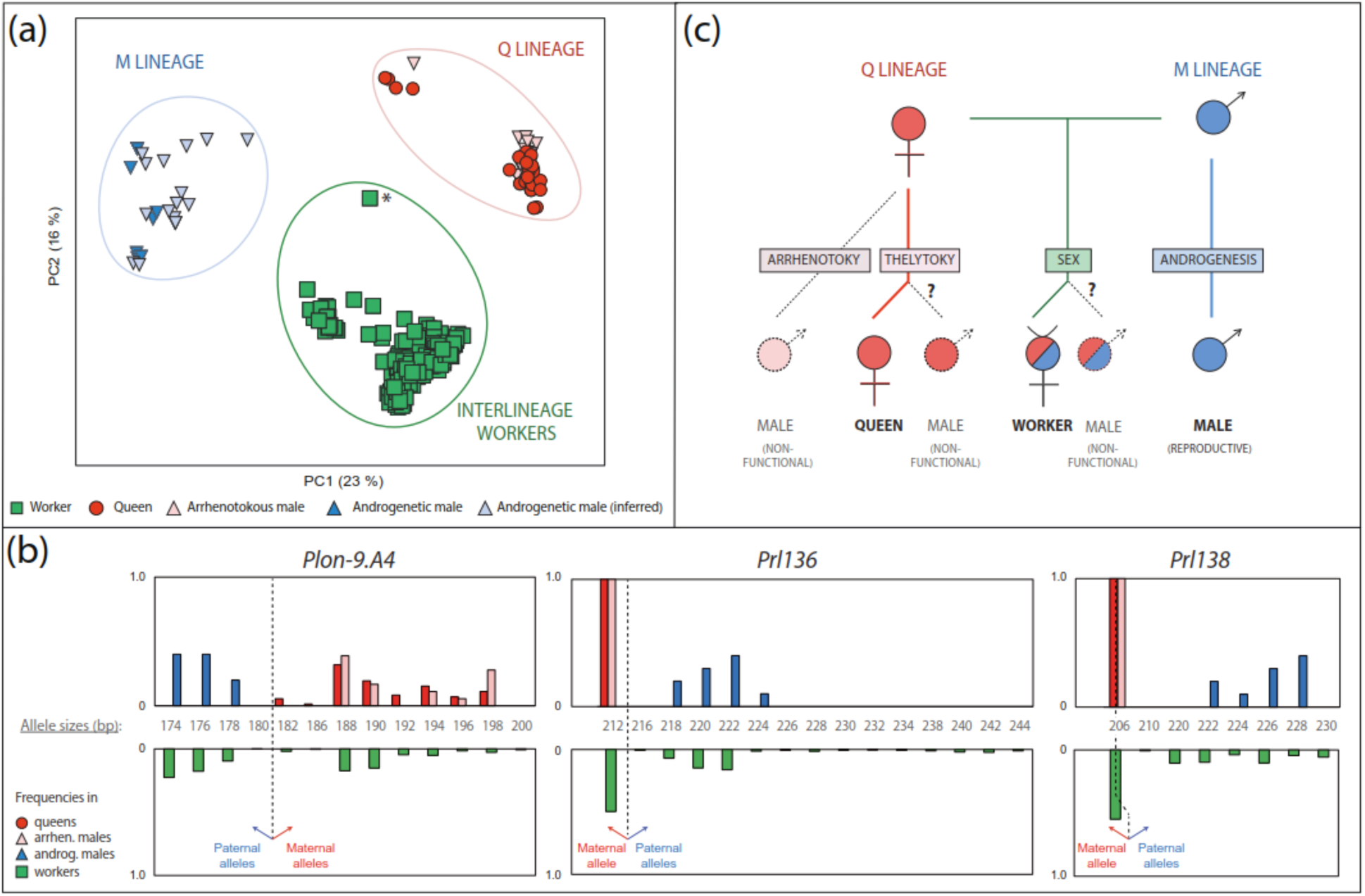
Two interdependent lineages (Q and M) coexist and interbreed for the production of workers across the range of *Paratrechina longicornis.* (a) Distribution of genetic diversity in queens, males and workers sampled in different localities across the world (see Fig. 1). The first two axes from a principal coordinates analysis (PCoA) based on microsatellite Bruvo’s distance are shown with the percentage of total variation explained by the axes. In localities with both queen and worker samples, male genotypes were inferred from workers. The worker indicated with a star was collected from the putative native range in Nepal and carried rare alleles. (b) Examples of how workers’ parental alleles are inferred. Three of the 19 microsatellite loci at which Q and M lineages had non-overlapping allele size ranges in the 22 localities where reproductive individuals were sampled are shown (other markers are shown in Fig. S1 and Fig. S2). Top panels: allele frequencies of Q lineage (N = 37 queens and N = 17 arrhenotokous males) and M lineage (N = 10 androgenetic males) at each of the 3 loci. Bottom panels: allele frequencies of workers (N = 341 individuals from 252 localities). The dashed lines represent the boundaries between the maternal and paternal alleles of workers, corresponding to Q and M lineages, respectively. Boundaries were inferred from the comparison of reproductive individuals’ genotypes and used to infer the parental origin of workers’ alleles in localities in which reproductive individuals were not collected. The allele size ranges of Q and M lineages overlap at the 206 bp allele for marker Prl138. (c) Schematic representation of the double-clonal system of *P. longicornis*. New queens belong to Q lineage and are produced by parthenogenesis (thelytoky). Different types of males coexist. The reproductives belong to M lineages and develop from male gametes with no maternal nuclear genome contribution (androgenesis). Non-functional males (dashed lines) carry alleles from Q lineage and develop from unfertilized eggs (arrhenotoky) or from diploid eggs (unknown mechanism). Queens are mated with males from the M lineage only and produced inter-lineage workers by sexual reproduction.

**Table 1.** Information about localities and sample sizes genotyped for Paratrechina longicornis. Numbers of longhorn crazy ant localities, numbers of genotyped males, queens and workers, and inferred male and queen haplotypes/genotypes for each surveyed geographic region.

| Region | Sampled localities | Genotyped individuals |  |  | Inferred genotypes |  |
| --- | --- | --- | --- | --- | --- | --- |
|  |  | Male | Queen | Worker | Male | Queen |
| Arabia | 3 | - | - | 4 | 3 | 3 |
| Caribbean | 19 | 2 | - | 30 | 17 | 30 |
| Central America | 1 | 2 | 1 | 4 | - | 4 |
| East Asia | 36 | 17 | 16 | 55 | 34 | 32 |
| Indian subcontinent | 35 | 4 | 6 | 50 | 35 | 41 |
| North America | 8 | 1 | 2 | 14 | 7 | 6 |
| Northeast Asia | 22 | - | - | 22 | 22 | 22 |
| Oceania | 25 | - | - | 26 | 25 | 26 |
| South America | 4 | - | - | 4 | 4 | 4 |
| Southeast Africa | 9 | - | 1 | 22 | 9 | 11 |
| Southeast Asia | 88 | 5 | 9 | 103 | 89 | 92 |
| Southern Europe | 4 | 1 | 1 | 3 | 3 | 3 |
| West Africa | 2 | 1 | 1 | 4 | 1 | 1 |
| Total | 256 | 33 | 37 | 341 | 249 | 275 |

Queen genotypes were consistent with clonal reproduction through apomixis or central fusion automixis as previously suggested (Pearcy et al., 2011). These two modes of parthenogenesis result in a partial decrease of heterozygosity at each generation due to mitotic or meiotic recombination events (Rabeling and Kronauer, 2013). Accordingly, queens were all fairly homozygous (mean individual heterozygosity ± SD = 16 ± 8%, range: 0-30%; N = 37 queens from 19 localities; Fig. S1). Genotype variation among matrilines could be assessed within 17 localities (Data S1_ClonalReproduction). In 13 localities, maternal genotypes were either completely identical or displayed minor allelic differences which could be explained by the occurrence of recombination events during parthenogenesis (Pearcy et al., 2006). In four localities, female genotypes were not consistent with a single clonal matriline, suggesting that different clonal matrilines coexisted locally or that some queens were produced by intra-lineage sexual reproduction.

Three types of male genotypes were observed (Fig. 2c). (*i*) Twenty-five males including all the males inferred from queen-worker comparisons carried one allele from M lineage at each diagnostic locus, suggesting that they were produced clonally by androgenetic reproduction (N = 20 localities). (*ii*) Eighteen males carried one allele from Q lineage at each locus (N= 5 localities). These males had variable genotypes at loci where the queens were heterozygous (N=12 males from 2 localities), suggesting that they developed from unfertilized eggs with random segregation of maternal alleles as is typical for most hymenopteran males (arrhenotokous parthenogenesis). (*iii*) Five males with more than one allele per marker were found in another 3 localities. It is unclear whether these males were diploid individuals that were homozygous at the complementary sex determination locus/loci, if any, or chimeras resulting from cytological incidents (Aamidor et al., 2018; Michez et al., 2009).

The production of each type of male was not confined to particular localities as multiple male types could be found within several localities (Data S2_Global_Genotypes). To gain insights into the relative frequency of each type, we genotyped males produced in two laboratory colonies. Out of 223 males, 80 males were inferred to be androgenetic, 127 were inferred to be arrhenotokous, and 16 had more than one allele per locus, suggesting that these were diploids or chimeras. To investigate whether arrhenotokous males are functional, we inspected their reproductive system and genotyped the spermathecal contents of queens. Live sperm was observed in seminal vesicles of arrhenotokous males (N=20), but all dissected queens carried sperm with a single allele from M lineage at each locus, suggesting that they had mated with androgenetic males only (N=86).

To summarize, workers sampled in 252 localities across four continents were first-generation hybrids of the same two nuclear lineages. The two lineages appeared to be maintained genetically distinct over generations through clonal reproduction of each sex: the genome of M lineage is transmitted through androgenesis from males to males. In contrast, the genome of Q lineage is parthenogenetically transmitted from queens to queens. Although males carrying Q alleles occur regularly, they do not seem to readily reproduce. Altogether, our results suggest that double-clonality is not a local phenomenon in *P. longicornis*.

### Clonal diversity

PCoA analyses of the queen and male genotypes from the 22 localities with reproductive individuals suggested the existence of multiple sub-groups within the queen and androgenetic male lineages (Fig. S5a and S5b). To further investigate the two interbreeding lineages’ variability, we reconstructed the maternal and paternal haplotypes of all workers from the 234 localities where reproductive individuals were not available (Table S1 and Supplementary Text).

The analysis of the 257 queen and 256 male genotypes across the 256 localities surveyed revealed that the allelic diversity of the M lineage was 2.7 times greater than that of the Q lineage (N = 253 genotypes per lineage, after reduction to one genotype per locality per lineage; Table S2).

**Table 2.**
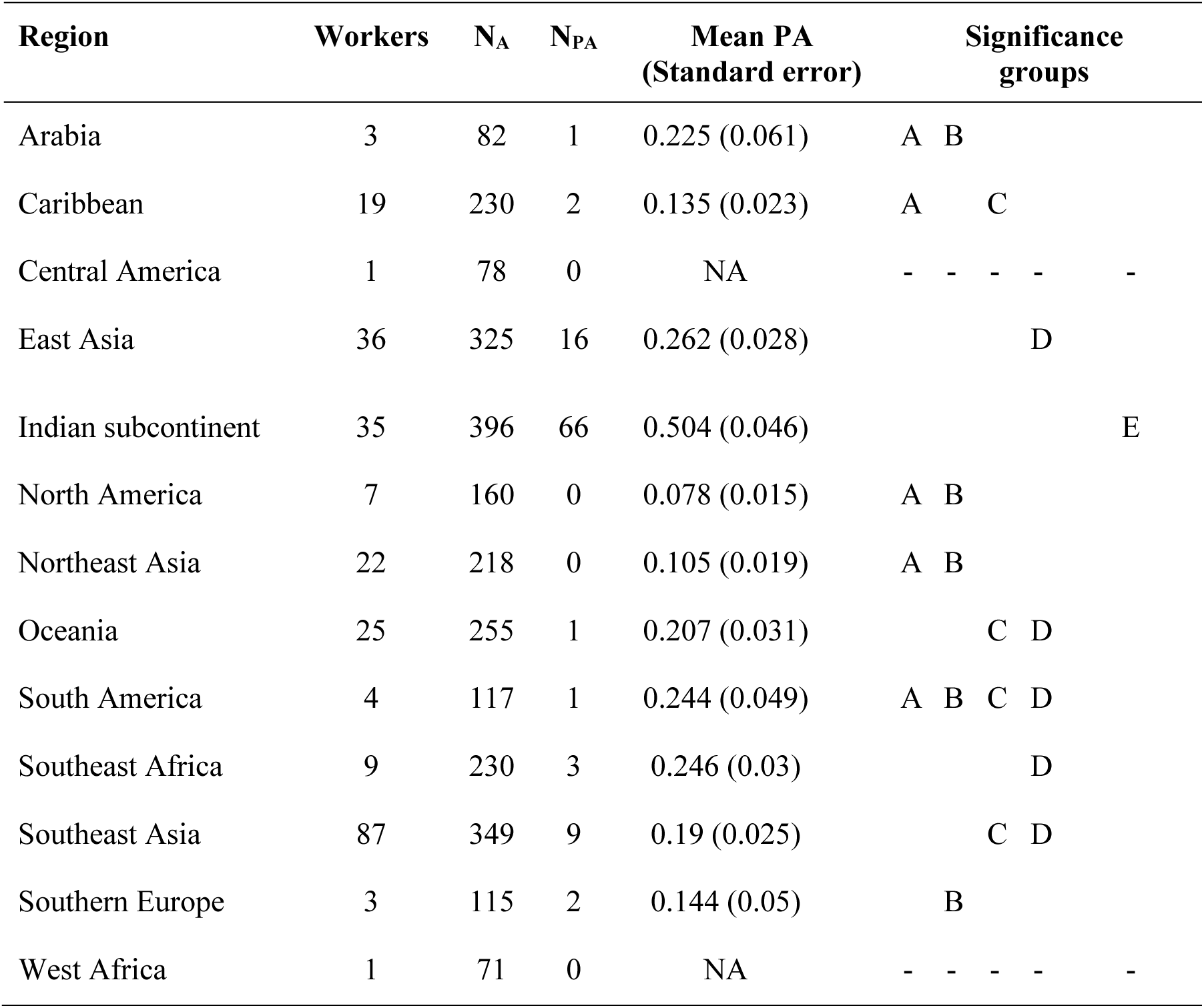
Distribution of private microsatellite alleles across the range of longhorn crazy ant. The number of worker genotypes, number of alleles (N_A_), total number of private alleles (N_PA_), and mean number of private alleles per locus (Mean PA) across the 40 microsatellite loci surveyed are indicated for each geographic region. The mean number of private alleles per locus was calculated using the rarefaction approach with a standardized sample size of 4. Differences in the mean number of private alleles among regions were analyzed using Kruskal-Wallis tests followed by post-hoc Wilcoxon signed-rank tests with a false discovery rate (FDR) correction. Regions that differ significantly (P< 0.05) in their mean private allele richness are indicated by different sets of letters.

| Region | Workers | $N_A$ | $N_{PA}$ | Mean PA<br>(Standard error) | Significance<br>groups |
| --- | --- | --- | --- | --- | --- |
| Arabia | 3 | 82 | 1 | 0.225 (0.061) | A B |
| Caribbean | 19 | 230 | 2 | 0.135 (0.023) | A C |
| Central America | 1 | 78 | 0 | NA | - - - - - |
| East Asia | 36 | 325 | 16 | 0.262 (0.028) | D |
| Indian subcontinent | 35 | 396 | 66 | 0.504 (0.046) | E |
| North America | 7 | 160 | 0 | 0.078 (0.015) | A B |
| Northeast Asia | 22 | 218 | 0 | 0.105 (0.019) | A B |
| Oceania | 25 | 255 | 1 | 0.207 (0.031) | C D |
| South America | 4 | 117 | 1 | 0.244 (0.049) | A B C D |
| Southeast Africa | 9 | 230 | 3 | 0.246 (0.03) | D |
| Southeast Asia | 87 | 349 | 9 | 0.19 (0.025) | C D |
| Southern Europe | 3 | 115 | 2 | 0.144 (0.05) | B |
| West Africa | 1 | 71 | 0 | NA | - - - - - |

PCoA (Fig. 3b) separated queens into two main groups (referred to as Clone A and Clone B) which differed at seven of the 16 loci analyzed (Fig. S6) and appeared widespread in invasive populations (Fig. 3a). Four inferred Nepalese queens carrying rare alleles had distinct PCoA coordinates and could not be assigned to a particular clone. Most queens belonged to Clone A (233 of 257 genotypes). These queens carried identical homozygous genotypes at 14 of the 16 loci analyzed, with only rare, seemingly recent, mutations in some individuals. The two remaining loci were more variable and occasionally heterozygous (*Plon-9.A4* was heterozygous in 30 of 32 queens sampled, while *Plon-8.F2* was heterozygous in 1 of 32 queens sampled and potentially heterozygous in queens of the pl381 and plvn07 localities, where nestmate workers carried different maternal alleles). Queens of Clone B were relatively less common (20 of 257 genotypes). Most variations within this second clone appeared to stem from recent mutations (one repeat mutation present in a small group of individuals).

**Fig. 3.**
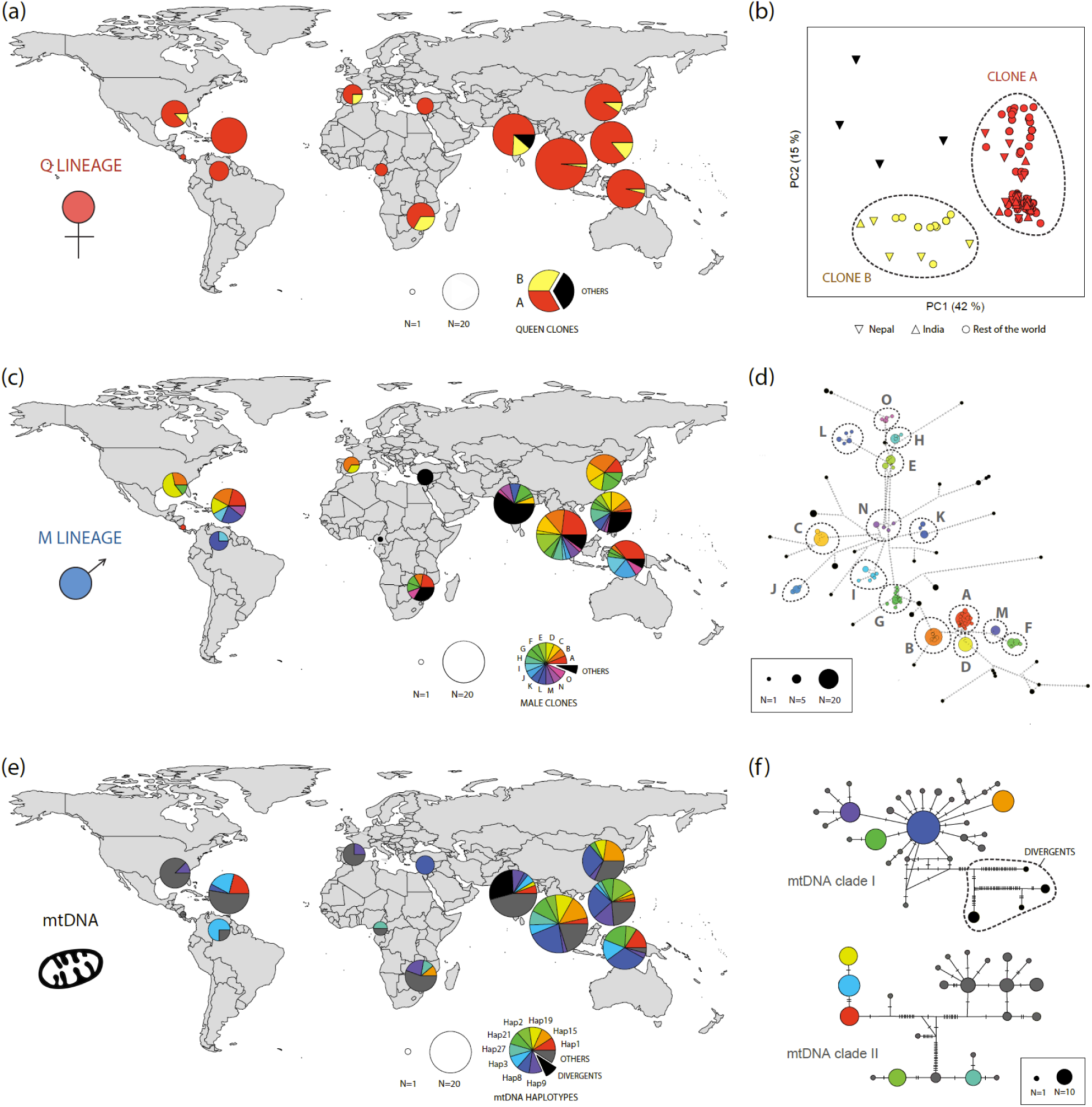
Phylogeography of *Paratrechina longicornis:* microsatellite variation of Q lineage (a-b) and M lineage (c-d), and mitochondrial DNA variation (e-f). For each panel pair, colors correspond to divergent genetic groups or haplotypes that are commonly encountered. Rare genotypes/haplotypes are depicted in black. The size of pie charts on the maps are functions of the sample sizes (ln(N)+0.5); (a, c, e). (a-b) Diversity of the Q lineage based on 257 microsatellite genotypes of queens (256 localities). (a) Frequencies of the 2 main clonal lineages identified through principal coordinate analyses. (b) First two axes from a principal coordinate analysis based on microsatellite Bruvo’s distance with the percentage of total variation explained by the axes. Genotypes from the putative native range (India and Nepal) are shown with different symbols to highlight their outlier positions. (c-d) Diversity of M lineage based on 256 microsatellite genotypes of androgenetic males (253 localities). (c) Frequencies of the 15 main clonal lineages identified by poppr across the 13 geographical regions surveyed. (d) Median-joining network of multilocus male haplotypes. Each haplotype is represented by a disk and tick marks on the lines joining haplotypes represent the number of mutated positions. Disk sizes are proportional to sample sizes. Disk colours and letters correspond to the multilocus clonal lineages of the A panel. (e-f) mtDNA variation based on 255 haplotype sequences of a mitochondrial region of 1,726 bp containing the COI and COII genes (254 localities). (e) Frequencies of the 9 most common mtDNA haplotypes, as well as of the 4 ’divergent’ haplotypes observed in the phylogenetic analyses (see panel f). (f) Median joining mtDNA haplotype network. Disk sizes are proportional to the number of sequences sampled for each haplotype. Single nucleotide substitutions are shown as ticks on the connecting lines between haplotypes. The haplotype network was divided into two divergent clades as discussed in Tseng et al. 2019 (see also Fig. SI7).

One hundred-thirty-eight different haploid genotypes were observed among the 256 androgenetic males analyzed. A median-joining network of the genotypes revealed a star-like structure with lower-frequency genotypes at the tips of the branches (Fig. 3d), consistent with clonal radiation (Weetman et al., 2002). Alternative genotypes differing by one or a few repeats were collapsed into 15 main multilocus clonal lineages (MLLs) accounting for 83% of the sampling. These MLLs appeared randomly distributed across the world (Fig. 3c), and this notion was supported by the fact that only 10 % of androgenetic male genetic variation could be explained by differences among geographical regions (AMOVA based on Bruvo’s distances, p<0.001, 9999 permutations; 215 males from 11 regions).

In *P. longicornis,* new males and queens mate within their natal nest and dispersion occurs primarily through colony budding (Passera, 1994). If no dispersion occurs among colonies, the queen and male genomes should evolve in parallel leading to an association between queen clones, maternal mtDNA and male MLLs. No strong correlation between the and male nuclear variations was observed and no one-to-one association between the 15 most common males’ MLLs and the 13 most common maternal mtDNA haplotypes was found. Yet, some male MLLs appeared to mate more frequently with one queen clone than the other (Table S3). For example, less than one percent of the queens of Clone A (2 out of 233) were found mated one of the four clones associated with queens of Clone B (N = 19 queens). Similarly, 19 % of male genetic variation could be explained by the difference among maternal mtDNA haplotypes (AMOVA based on Bruvo’s distances, p<0.001, 9999 permutations; 217 males carrying 22 haplotypes). Altogether, these results suggest that the queen and male genotypes are not totally randomly assorted, but that movements of reproductive individuals among colonies may erase patterns of parallel clonal evolution. A candidate for such gene flow could be the occasional fusion of genetically distinct colonies following multiple introduction events as seem to have happen in one of our sampled localities where four different androgenetic clones, probably brought together with plant materials were observed in the same flowerbed (locality plID04).

### Variation in mtDNA

We identified 55 mtDNA haplotypes. Of these, 15 were common haplotypes found in more than one geographic region and appeared to be randomly distributed (Fig. 3e-f). The two queen clones did not share any mtDNA haplotypes (46 haplotypes were observed in localities headed by queens of Clone A, while 6 haplotypes were observed in localities with queens of Clone B). Despite the haplotype segregation following the clone origin (Fig. 3f), the patterns of queens’ nuclear and mitochondrial variations were not congruent. For example, identical queen genotypes were regularly associated with mtDNA haplotypes originating from the divergent mtDNA clades identified from previous studies (Tseng et al., 2019b). The average genetic distance between these two mtDNA clades was 5.7%, translating into 1.6 to 2.5 million years of divergence using standard mtDNA evolutionary rates of 2.3 to 3.54% My^-1^ (Papadopoulou et al., 2010). If queens’ mitochondrial and nuclear genomes had linked histories, such large divergence time would have resulted in significant microsatellite genotypic differences given that microsatellites have a typical mutation rate of about 10^−4^ mutations per locus per generation (Crozier et al., 1999; Weber and Wong, 1993). These results suggest the occurrence of past introgression events that broke the physical linkage between the nuclear and mitochondrial genomes of queens.

In most organisms, mtDNA is maternally deposited. To determine if androgenetic males also inherit mtDNA from their mother, we compared the mtDNA haplotypes of androgenetic males, queens and workers in nine localities. Analyses indicated that all castes shared the same haplotype within each of the localities studied consistent with a shared mode of maternal mitochondrial DNA transmission among castes (Data S3_colony-mthaplotype). These results suggest that androgenetic males are cytonuclear hybrids carrying paternal nuclear DNA and maternal mtDNA.

### Inference of the native range

Analyses based on reproductive individuals’ genotypes at 16 microsatellite loci suggested the occurrence of rare clonal genotypes in individuals collected from the Indian subcontinent (see Fig. 3a and 3c). To explore further whether this region is a genetic diversity hotspot for the species, we compared the complete genotypes of workers at 40 microsatellite loci. We identified 101 private alleles (ranging from 0 to 66 per geographic region) in the 252 workers sampled worldwide. The mean number of private alleles was significantly higher in the Indian subcontinent (65%) than in any other geographic region although it accounted for only 14 % of the sampling (Table 2).

Analysis for mtDNA diversity showed that 40 of the 55 different haplotypes observed were private haplotypes to one geographic region (Fig. S7). Southern Europe had the highest frequency of private haplotypes (75%; 3 of 4 sequences), followed by the Indian subcontinent (60%; 21 of 35 sequences) and Southeast Africa (56%; 5 of 9 sequences; Table S4). However, the private mtDNA haplotypes from Southern Europe and Southeast Africa were all closely related to widespread haplotypes (divergence of 25,190 to 50,380 years using a conservative low divergence rate of 2.3 % My^-1^) (Papadopoulou et al., 2010), suggesting these private haplotypes likely result from recent mutations of common invasive haplotypes. By contrast, sequences with no close relationship to any widespread haplotypes were only found in the Indian subcontinent. More specifically, four Himalayan haplotypes (Hap31, Hap37, Han 46, and Hap48) appeared unrelated to any other haplotypes sampled in our study (minimum divergence of 378,261 to 1,008,696 years; Fig. 3f). The private haplotypes found in the Indian subcontinent are suggestive of the diverse gene pool of a native range. Rarefaction analyses indicated that the Indian subcontinent harbored the highest haplotype/allele richness for both mtDNA (Fig. 4a) and microsatellites (Fig 4b). Therefore, our results suggest that the Indian subcontinent is a genetic diversity hotspot for *P. longicornis* and likely constitutes the native range of the species.

**Fig. 4.**
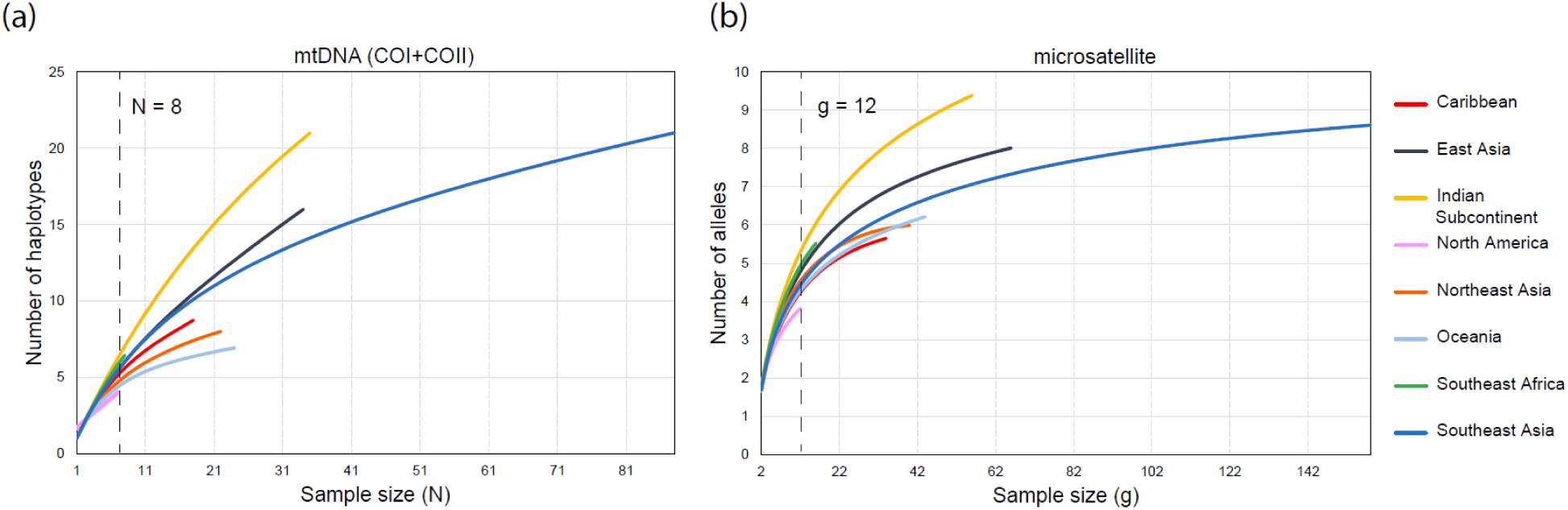
Rarefaction curves of (a) mtDNA haplotype richness (number of distinct mtDNA haplotypes) and (b) microsatellite allele richness (mean number of distinct alleles per locus) of *Paratrechina longicornis* workers for each geographic region. Sample sizes correspond to the number of mtDNA sequences sampled (N) and the number of microsatellite alleles sampled per locus (g; 2 per individual). South America, Arabia, and Southern Europe were not included in these analyses due to small sample sizes (N < 5; g < 10). The vertical dashed lines indicate the lowest sample size in any of the remaining regions.

## DISCUSSION

Our analyses of genetic variation across the range of the invasive ant *P. longicornis* reveal that this cosmopolitan species most likely originate from the Indian subcontinent, and that all populations of this species surveyed in both its invasive and inferred native ranges display a double-clonal system with the presence of two divergent, yet interdependent queen and male clonal lineages. Double-clonality was initially reported in a single Thailand population of *P. longicornis* (Pearcy et al., 2011). This study extends this finding and shows that this genetic system is widespread across the species range. Queens appear to be typically produced by thelytokous parthenogenesis and inherit maternal DNA only, while functional males are androgenetic cytonuclear hybrids carrying paternal nuclear DNA and maternal mitochondrial DNA. Workers are always highly heterozygous individuals originating from crosses between the divergent queen and male lineages. Remarkably, our phylogeographic analyses revealed that all clonal queens and clonal males of *P. longicornis* belong to two respective, non-recombining Q and M lineages that have remained divergent across time and space. This result was supported by the segregation of queen and male samples from different continents in two divergent groups (22 localities) and the finding that workers always carry F1 hybrid combinations of these two parental groups at multiple diagnostic markers (256 localities). Interestingly, M lineage appears far more variable than Q lineage. This could be partially explained by different mutation rates among the two *P. longicornis* lineages that result from differences in ploidy: Q lineage is carried by diploid individuals which undergo a modified meiosis, while M lineage is only found in the haploid stage and never goes through meiosis (Sharp et al., 2018). Alternatively, selection may be much stronger in queens than males, leading to stronger clonal sweeps in the former. Ant males are short-lived and their only task is to mate, which is expected to limit the opportunity for selection (Feldhaar et al., 2008).

To date, double-clonal reproduction has been reported in another three ant species, *Wasmannia auropunctata* (Fournier et al., 2005), *Vollenhovia emeryi* (Kobayashi et al., 2008; Ohkawara et al., 2006) and *Cardiocondyla kagutsuchi* (Okita and Tsuchida, 2016). In contrast with *P. longicornis,* recombination among lineages occurs in at least one of these species, *W. auropunctata,* through the production of functional interlineage hybrid queens by sexual reproduction (Foucaud et al., 2007). In *V. emeryi*, interlineage hybrid queens are also observed in both field and laboratory colonies, but whether these events lead to recombination among lineages over evolutionary time remains to be determined (Miyakawa and Mikheyev, 2015; Okamoto et al., 2015). The complete segregation of *P. longicornis* parental genomes in the two lineages is reminiscent of social hybridogenesis, another genetic system which has been reported in the ant genera *Pogonomyrmex*, *Solenopsis*, *Messor* and *Cataglyphis*. Under this system, two lineages consisting of both queens and males coexist across populations and interbreed regularly for the production of workers (Lacy et al., 2019; Leniaud et al., 2012; Norman et al., 2016; Romiguier et al., 2017; Sirviö et al., 2011). Social hybridogenesis has been shown to be maintained by genetic caste determination systems, whereby the fate of a female egg is determined by its genotype, rather than by environmental factors as is usually the case in social Hymenopterans (Cahan et al., 2004; Darras et al., 2014; Schwander et al., 2010). A similar mechanism likely underlies the maintenance of double-clonality in *P. longicornis*.

One unexpected finding of our study is the frequent occurrence of arrhenotokous males developing from unfertilized eggs. Arrhenotokous males were collected in five localities and were also regularly produced in laboratory colonies. Dissections of males’ and queens’ reproductive organs indicated that these arrhenotokous males could produce sperm, but virtually never mate with queens. Yet, rare reproduction events involving arrhenotokous males may have played a key role in the spread of the double-clonal system. The comparison of nuclear and mitochondrial DNA variations suggested that the nuclear genome of clonal queens has introgressed into a diverse mitochondrial background leading to cyto-nuclear incongruences. A candidate mechanism for such introgression could be reproduction by arrhenotokous males. Following the emergence of clonality, clonality-inducing mutation(s) may have spread contagiously to neighboring populations through arrhenotokous males carrying the mutation(s)(Simon et al., 2003). That fertilization by the Q lineage sperm of an arrhenotokous male may occasionally lead to queen development is also interesting. As mentioned above, the perfect association between the genotype of females and their caste indeed suggests the existence of a hardwired genetic caste determination system, whereby diploid eggs carrying Q genome only develop into new queens. Fertilization of a Q lineage egg by the Q lineage sperm of an arrhenotokous male should therefore lead to queen development. Such sexual production of new queens, albeit rare, would allow the purging of deleterious mutations and the generation of new allelic combinations through recombination. Occasional recombination events have been reported in other clonal ants such as *Mycocepurus smithii* (Rabeling et al., 2011), *Oocerea biroi* (Trible et al., 2020) or desert ants of the genus *Cataglyphis* (Darras et al., 2014; Doums et al., 2013).

The patterns of both nuclear and mitochondrial variations placed the genetic diversity hotspot of *P. longicornis* in the Indian subcontinent. Rare clonal lineages were also identified only in this region, consistent with this geographical region being part of the native range of the species. Surprisingly, however, the three other species of the *Paratrechina* genus (*P. ankarana, P. antsingy* and *P. zanjensis*) are all endemic to Africa (Boudinot et al., 2021; LaPolla and Fisher, 2014)*. Paratrechina umbra,* a species known only from China, was recently included in the *Paratrechina* genus (Williams and Lapolla, 2016), but phylogenetic analyses indicate that this species belongs to the genus *Nylanderia* (unpublished data). The incongruence between the inferred South Asian origin of *P. longicornis* and the African distribution of its sister species may originate from an intercontinental dispersal event which resulted in a disjunctive Afro-Asian distribution of the *Paratrechina* species, similar to other ant genera such as *Bothroponera* and *Parasyscia* (Borowiec, 2016; Joma and Mackay, 2020).

The global spread of ant species has been greatly facilitated by major events along recent human history, especially during the last two waves of globalization (1850–1914 and 1960–today) (Bertelsmeier et al., 2017). The distribution of *P. longicornis* rapidly expanding after 1860 coincides with the first wave of globalization (Bertelsmeier et al., 2017). In the late 19th century, the domestic and foreign trade of India grew exponentially (Kuehlwein, 2021), and by 1900, India had become the largest exporter in Asia and ranked the ninth largest in the world (Hanson, 1980; Kuehlwein, 2021). Such India-outbound commerce activities may have fueled the early trans- continent spread of *P. longicornis* (i.e., from its native range to regions across the world). Furthermore, our findings of multiple, randomly distributed clonal sub-lineages and mtDNA haplotypes within each region of the species’ invasive range suggests that multiple introduction events involving genetically distinct propagules seems common. Border inspection records worldwide are consistent to our interpretation of repeated, multi-origin introductions of this ant around the world as *P. longicornis* is one of the most frequently intercepted species (Bertelsmeier et al., 2018; Lee et al., 2020; Ward et al., 2006).

In addition to the high degree of propagule pressure, *P. longicornis* possesses several traits that may have contributed to its worldwide spread. It forms colonies with hundreds of reproductive queens and is highly adapted to disturbed environments (Wetterer, 2008). More importantly, its double-clonal reproductive system eliminates the risk of inbreeding depression during colonization as workers are always offspring of queens and males carrying divergent genomes even when these reproductive are siblings (Pearcy et al., 2011; Trager, 1984). The current study shows that double- clonal reproduction is present in the inferred native range of the species, indicating that this reproduction system may have pre-adapted *P. longicornis* to be a successful invader. A similar case can be observed in another highly invasive ant, the little fire ant, *W. auropunctata*: while sexual and double-clonal populations are both observed in the ant’s native range, all invasive populations are double-clonal (Foucaud et al., 2009, 2007; Fournier et al., 2005). We argue that both reproduction modes may also coexist in the native range of *P. longicornis* but remain undetected in our sampling. Sexual populations of *W. auropunctata* generally occurred in natural habitats, while clonal populations are more frequent in disturbed habitats (Foucaud et al., 2009). Since our sampling of *P. longicornis* mostly focused on human-disturbed areas, a comprehensive sampling regime across the native range that includes both disturbed and undisturbed habitats would hold the key to elucidate whether native sexual populations exist in *P. longicornis* and how double-clonality emerged.

## MATERIALS AND METHODS

### Sample collection

We collected *P. longicornis* in 67 localities and performed additional genetic analyses on previously collected samples from 189 localities (Tseng et al., 2019a, 2019b). In total, we obtained samples from 256 different localities across the range of the species (32 from America, 11 from Africa, 4 from Europe, 184 from Asia and 25 from Oceania; Fig. 1). To study the distribution of queen and male clonality, reproductive individuals were sampled in 22 representative localities, including the original locality in Thailand where double-clonality was the first discovered (Pearcy et al., 2011) (Table 1).

Our sampling revealed three different types of males (see Results). To investigate the evolutionary significance of these males, two colony fragments consisting of ∼50 reproductive queens and a few thousand workers were collected in two neighboring Spanish populations (Malaga) and maintained in laboratory conditions. Some of the emerging males were randomly dissected to assess whether they produced sperm and were preserved for genetic analyses. The remaining males were left to mate with new queens within the nests. After a year, the sperm storages of the new generation of queens were collected to gain insight into the mating success of each male type (see *Distribution and characteristics of double-clonality* for the detailed methodology).

### Microsatellite genotyping

Queens, males, and a sample of workers (mean ± SD = 4.1 ± 3.0 per locality) were genotyped in 22 localities. We also analyzed 1-2 workers for each of the remaining 234 localities. A total of 411 selected individuals were genotyped at 40 microsatellite loci using published primer sequences (MERPDC et al., 2011; Tseng et al., 2019a); Data S4_List_of_microsatellite_primers). DNA extraction and genotyping were carried out as described in Tseng, Darras et al. 2019a. For some localities, partial genotypes were released earlier to answer other research questions (Tseng et al., 2019a, 2019b).

Queen, male and sperm samples collected from the two laboratory colonies were genotyped at 13 microsatellite loci (see Data S4_List_of_microsatellite_primers). Microsatellite data derived from these colonies were analyzed separately from the main dataset. All genotypes are available as supporting information (Data S2_Global_Genotypes and Data S5_Laboratory_genotypes).

### Sequencing and processing of mtDNA

To compare nuclear and mitochondrial variation patterns, a portion of the mitochondrial genome containing the *cytochrome oxidase subunit I*, *cytochrome oxidase subunit II* and *tRNA–Leu* genes, and an intergenic spacer was sequenced and analyzed following methods described by Tseng *et al*. 2019b. A female from each of the 67 new localities sampled here was sequenced (63 workers and 4 queens). Additional castes were sequenced in 9 localities (both sexes were sequenced in 4 localities and one individual of each caste was sequenced in each of the remaining 5 localities). These sequences were aligned with published sequences of *P. longicornis* (GenBank accession numbers: KY769964-KY770017). The region containing the intergenic spacer and the *tRNA–Leu* gene showed a low level of conservation and was removed from the alignment. For further analyses, the sequences were collapsed into haplotypes using DnaSP v5.10 (Librado and Rozas, 2009). The new haplotype sequences identified in this study were deposited in GenBank (MZ398110- MZ398127).

### Distribution and characteristics of double-clonality

The double-clonal mode of reproduction previously described in Thailand resulted in different genetic makeups for each caste: queens and males belong to different genetic groups, while workers are highly heterozygous individuals carrying alleles from both groups at each marker. To investigate the global distribution of double-clonality, we compared the microsatellite genotypes of reproductive individuals and workers (22 and 252 localities, respectively). Workers’ paternal genotypes were reconstructed based on the genotypes of mothers when the latter was available (Pearcy et al., 2011). To enable comparative analyses of male and female genotypes, the genotypes of haploid males were encoded as diploids by doubling their alleles. Allelic frequencies and individual heterozygosity (i.e., number of heterozygous loci divided by a total number of loci analyzed per individual) were estimated using GENALEX v6.502 (Peakall and Smouse, 2012). *Fij* kinship coefficients were estimated according to (Loiselle et al., 1995) using SPAGeDi v1.5 (Hardy and Vekemans, 2002). Genetic variation was investigated using principal-coordinate analyses (PCoA) based on genetic distances. Distances were estimated with the R package ’poppr v2.9.1’ (Kamvar et al., 2014) using the Bruvo’s method, which considers mutational distances between microsatellite alleles (Bruvo et al., 2004). PCoAs were then performed on raw distances using GENALEX. The Bayesian clustering method implemented in STRUCTURE v2.3.3 was also used to determine the number of genetic lineages (K) among reproductive individuals (Pritchard et al., 2000). Only one queen and one male per locality were included in this analysis to avoid bias. The program was run 10 times for each value of K ranging from 1 to 10, with 200,000 Markov chain Monte Carlo iterations and a burn-in period of 50,000. Analyses were performed under the admixture model with correlated allele frequencies and without prior population information. The most likely genetic lineage numbers were determined using the ad-hoc Δ-K method (Evanno et al., 2005). The occurrence of clonal reproduction was assessed by comparing the genotypes of nestmate reproductive individuals when multiple queens or males were sampled within a locality. In addition, to gain insight into the ubiquity and reproductive ability of different types of males, laboratory-produced males, and the sperm storages of reproductive queens from the same colonies were genotyped (Data S5_Laboratory_genotypes).

Clonal (androgenetic) males inherit no nuclear material from their mother (Schwander and Oldroyd, 2016). To determine whether androgenetic *P. longicornis* males still inherit mitochondrial DNA from their mother, we compared the mtDNA sequences of male and female nestmates in 7 localities.

### Clonal diversity

Our results indicate that queens and males of *P. longicornis* belong to two widespread, non-recombining lineages, respectively (see Results). To analyze the distribution of genetic diversity within each of these two lineages, two (queen and male datasets) sub-datasets were reconstructed, each consisting of the genotypes of reproductive individuals sampled in the field (22 localities), as well as parental haplotypes inferred from workers (250 localities, including 16 of the 22 localities mentioned before). Sixteen loci had virtually no overlap between the allele size ranges of the clonal queen and male samples (Table S1). Microsatellite evolution typically follows a stepwise mutation model whereby allele size changes by a single repeat unit per mutation (Ellegren, 2004). This process generates unimodal allele size distributions with single repeat increments between successive alleles. If any, unsampled queen and male alleles were expected to be close in size to those observed in our reproductive individuals. This allowed assigning the parental origin of a worker’s allele at each of these 16 diagnostic loci solely based on its size when reproductive individuals were not available for deductive inferences (see Supplementary Text). The 24 remaining loci had overlapping allele size ranges between the two lineages and were not considered for this analysis. When multiple workers were sampled in a locality, redundant inferred parental genotypes, if any, were discarded.

To study genetic variation in clonal (androgenetic) males, we constructed a median-joining haplotype network of microsatellite genotypes using Network Fluxus v10.2 with the epsilon parameter set to 0 (Bandelt et al., 1999). This approach requires that microsatellite loci are linked, which we assumed to be the case as haploid male genomes are transmitted as complete linkage groups during androgenesis. Male haplotypes were clustered into multilocus clonal lineages (MLL) using a Bruvo’s distance threshold of 0.2 with the *mlg.filter* function of the R package ‘poppr v2.9.1’. To study queen genetic variation, we performed PCoA analyses instead as queens have diploid genomes that may recombine during parthenogenesis (Pearcy et al., 2006).

The queen and male datasets were reduced to one genotype per locality per lineage to compare diversities of the two lineages. Analyses of molecular variance (AMOVA) were performed with GENALEX using Bruvo’s distances.

### Inference of the native range

To infer the native range of *P. longicornis*, we examined whether some parts of the world displayed higher microsatellite and mtDNA richness than the rest of the species range. We assumed that genetic diversity would be lower in invasive populations due to genetic bottlenecks during colonization (Caron et al., 2014). We only used one mtDNA sequence per locality and one worker genotype (taking the first worker genotyped) per locality to avoid biases. In addition, indexes were corrected for differences in sample sizes among geographical regions using rarefaction methods. Microsatellite allelic richness and mean number of private alleles (e.g., alleles restricted to one geographical region) were estimated with the rarefaction algorithm implemented in ADZE v1.0 (Szpiech et al., 2008), while mtDNA haplotype richness was estimated using the rarefaction method of PAST v2.17c (Hammer et al., 2001). To visualize the geographical distribution of private mtDNA haplotypes, a median-joining mtDNA haplotype network was constructed using POPART (Leigh and Bryant, 2015).

## Acknowledgements

Morgan Pearcy performed preliminary phylogeographic analyses. We thank Amy Low, Ni-Chen Chang, Chung-Wei You, Chùn- Têng Coody Chiu, Chun-Yi Lin, Ching-Chen Lee, En-Cheng Yang, Han-Chih Ho, Hui-Siang Tee, Hung-Wei Hsu, Jia-Wei Tay, Kean Teik Koay, Li Yan Gan, Mark Ooi, Nellie Wong, Peter G. Hawkes, Ping-Chih Lin, Singham Veera, Su-Chart Lee, Sylvain Hugel, Yi-Ming Weng, Yueh-Hua Wu, Yu-Fang Tseng, and Zhengwei Jong for assistance in field collection and Qiaowei M. Pan for comments on the manuscript.

## Additional information

### Author contributions

C.C.S.Y., S.-P.T., and H.D. originally conceived the ideas. J.W., L.K., and C.-Y.L. helped develop an overarching research programme and collect the samples. C.C.S.Y., L.K. and T.Y. applied for the grant. S.-P.T., H.D. and P.-W.H. performed the experiments. S.-P.T. and H.D. carried out the analyses and drafted the manuscript. All authors contributed substantially to editing the manuscript and approved the submitted version.

### Data availability

All data are available in the main text or the supplementary materials.

